# Hippocampal ensembles represent sequential relationships among discrete nonspatial events

**DOI:** 10.1101/840199

**Authors:** Babak Shahbaba, Lingge Li, Forest Agostinelli, Mansi Saraf, Gabriel A. Elias, Pierre Baldi, Norbert J. Fortin

**Affiliations:** Department of Statistics, University of California, Irvine, CA, USA; Department of Computer Science, University of California, Irvine, CA, USA; Department of Neurobiology & Behavior, University of California, Irvine, CA, USA

## Abstract

The hippocampus is critical to the temporal organization of our experiences, including the ability to remember past event sequences and predict future ones. Although this fundamental capacity is conserved across modalities and species, its underlying neuronal mechanisms remain poorly understood. Here we recorded hippocampal ensemble activity as rats remembered a sequence of nonspatial events (5 odor presentations unfolding over several seconds), using a task with established parallels in humans. Using novel statistical methods and deep learning techniques, we then identified new forms of sequential organization in hippocampal activity linked with task performance. We discovered that sequential firing fields (“time cells”) provided temporal information within and across events in the sequence, and that distinct types of task-critical information (stimulus identity, temporal order, and trial outcome) were also sequentially differentiated within event presentations. Finally, as previously only observed with spatial information, we report that the representations of past, present and future events were sequentially activated within individual event presentations, and that these sequential representations could be compressed within an individual theta cycle. These findings strongly suggest that a fundamental function of the hippocampal network is to encode and preserve the sequential order of experiences, and use these representations to generate predictions to inform decision-making.

## INTRODUCTION

In humans, the hippocampus is known to play a key role in the temporal organization of memory and behavior. This includes our ability to remember when past experiences occurred^1–4^, but also extends to our ability to use information about past experiences to imagine or predict future outcomes^5, 6^. Considerable research indicates that this capacity is conserved across species and applies across stimulus modalities^3, 4, 6^, yet its underlying neural mechanisms remain poorly understood. The emerging conceptual framework suggests that the propensity of the hippocampal network to generate and preserve sequential patterns of activity underlies this fundamental capacity^4, 6–10^, a view supported by two main lines of electrophysiological evidence. First, hippocampal ensemble activity tends to exhibit sequential firing fields during the presentation of individual stimuli or inter-stimulus intervals (also known as “time cell” activity)^11, 12^, which has been shown to provide a strong temporal signal within such task events^13, 14^. Second, hippocampal neurons have been shown to code for sequences of spatial locations under different experimental conditions^6, 15, 16^. Of particular interest here is evidence that hippocampal activity can represent sequences of past, current, and upcoming locations when animals run on a maze^17–20^ or pause at a decision point (vicarious trial-and-errors)^21^, conditions in which the hippocampal network displays prominent theta oscillations and is thought to be engaged in online processing of upcoming decisions and goals^6, 9^. However, in order to elucidate the hippocampal coding properties supporting our fundamental ability to remember and predict event sequences, it is critical to: (1) record neural activity under broader experimental conditions more characteristic of episodic memory in humans, such as across sequences of discrete nonspatial events unfolding across several seconds, and (2) directly link these sequence coding properties with correct sequence memory judgments.

To address this important issue, we recorded hippocampal ensemble activity as rats performed a challenging nonspatial sequence memory task with established behavioral and neural parallels in humans^22–24^. Taking advantage of this unique behavioral approach, we began by examining how the organization of sequential firing fields varied with stimulus identity and across the full sequence of non-overlapping events. We then developed statistical tools, including deep learning approaches, to reveal the sequential structure in which the representations of different types of information varied within individual stimulus presentations (i.e., within 1.2 seconds). More specifically, we used a latent representation learning approach to quantify the differentiation of each type of task-critical information (stimulus, temporal order, and trial outcome information), and a neural decoding approach to quantify the decoding probability of each stimulus in the sequence. Finally, using a simpler decoding model with higher temporal resolution, we tested whether the representation of past, present and future stimuli (which are normally experienced across several seconds) could be compressed within a single theta cycle. We present compelling evidence that this work, which integrates experimental and analytical approaches specifically developed to address this fundamental problem, provides novel insights into the function of the hippocampus.

## RESULTS

We trained rats to perform a nonspatial sequence memory task, which shows strong behavioral correspondence in rats and humans^22^ and depends on comparable circuits across species^23,24^. In rats, this hippocampus-dependent task involves repeated presentations of sequences of nonspatial stimuli (odors ABCDE) in the same odor port, and requires subjects to determine whether each item is presented “in sequence” (InSeq; e.g., AB<u>C</u>…) or “out of sequence” (OutSeq; e.g., AB<u>D</u>…) to receive a water reward (Fig 1). After reaching criterion on the task, animals were implanted with a microdrive and, over a few weeks, tetrodes were gradually lowered into the pyramidal cell layer of the dorsal CA1 region of the hippocampus. Neural activity (spikes and local field potentials; LFP) was then recorded as animals performed the task (Fig 2a; see^25^ for a detailed analysis of single-cell and LFP activity from this dataset). Here we focus on elucidating the temporal dynamics of CA1 ensemble activity supporting the memory for sequences of events.

**Figure 1.**
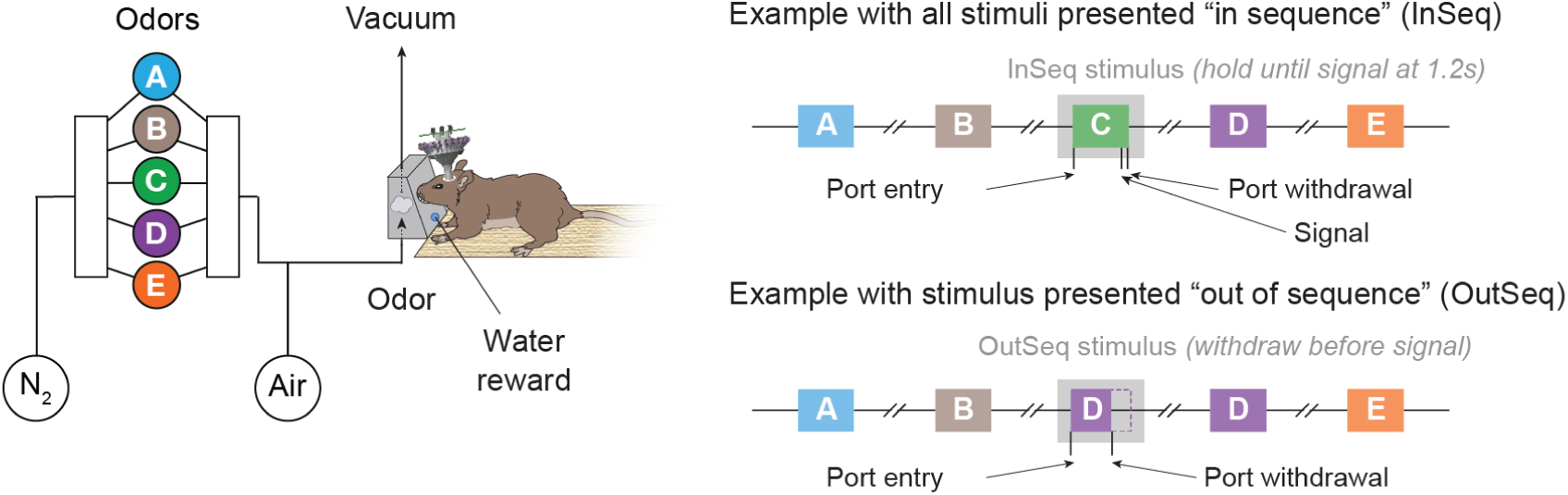
Nonspatial sequence memory task. Neural activity was recorded as rats performed a hippocampus-dependent sequence memory task with strong behavioral parallels in rats and humans^22^. The task involves repeated presentations of sequences of nonspatial stimuli and requires subjects to determine whether each stimulus is presented “in sequence” (InSeq; e.g., AB<u>C</u>…) or “out of sequence” (OutSeq; e.g., AB<u>D</u>…). Using an automated odor delivery system (*left*), self-paced sequences of five odors were presented in the same odor port (median interval between consecutive odors ∼5 s). In each session, the same sequence was presented multiple times (*right*), with approximately half the presentations including all stimuli InSeq (*top*) and the other half including one stimulus OutSeq (*bottom*). Each odor presentation was initiated by a nosepoke and rats were required to correctly identify the odor as either InSeq (by holding their nosepoke response until the signal at 1.2 s) or OutSeq (by withdrawing their nosepoke before the signal; <1.2 s) to receive a water reward.

**Figure 2.**
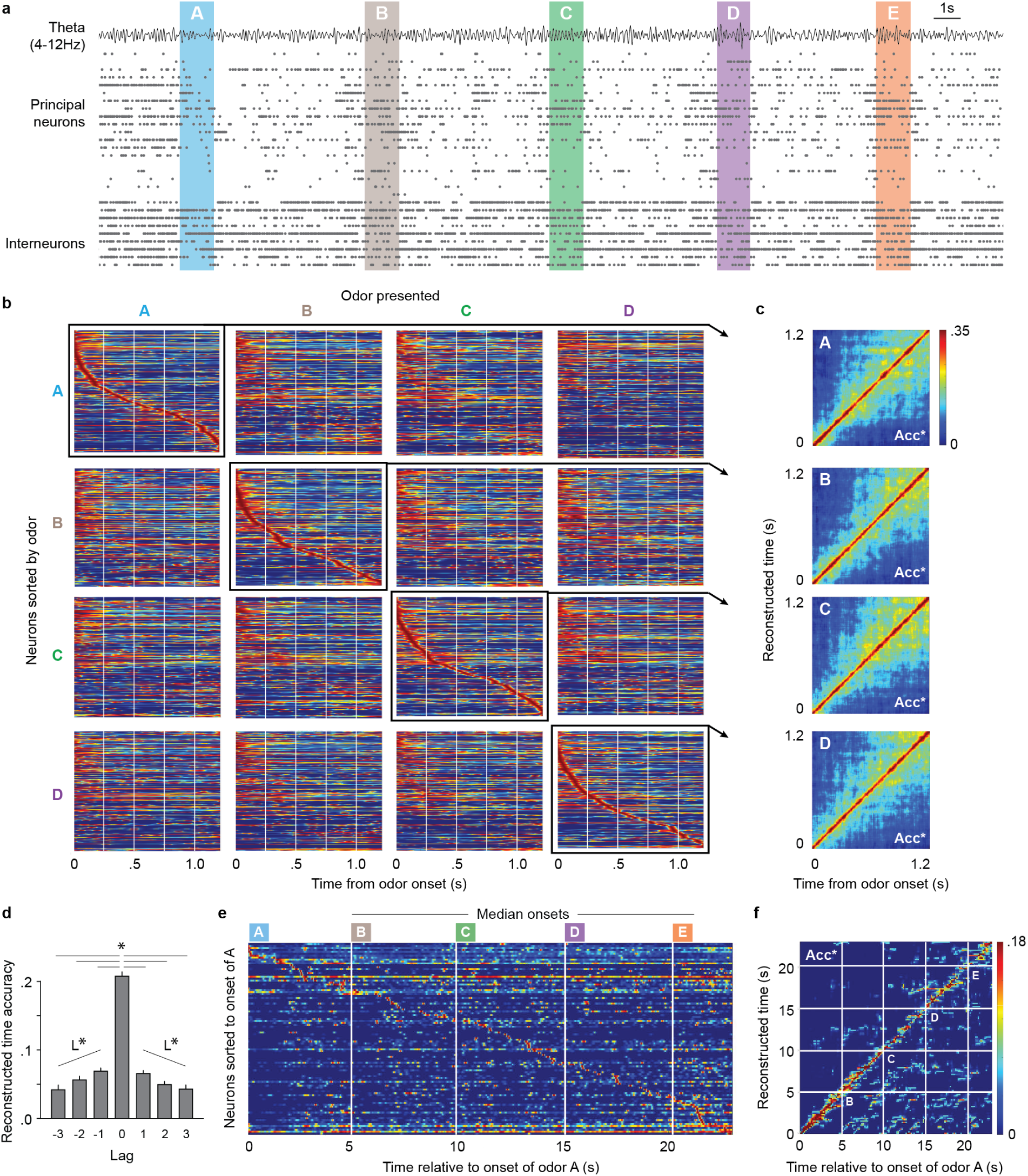
CA1 ensemble activity is organized in sequences of firing fields within individual stimuli and across the sequence. **a**, Example ensemble activity (putative principal neurons and interneurons) and theta oscillations (bandpass: 4-12 Hz) from representative subject during one sequence presentation. Neurons were sorted by their peak firing time in relation to the onset of odor A. **b**, The sequential organization of firing fields showed stimulus selectivity. Each peri-stimulus time histogram (PSTH; 150ms gaussian) shows the normalized firing rate of all active neurons during presentation of a specific odor type (red indicates the maximum firing rate of each neuron, and blue indicates 0 Hz; 267 active neurons for odor A, 192 for B, 209 for C, and 223 for D; neurons collapsed across five subjects, one session per subject). To visualize how sequential firing fields varied across odor types, PSTHs are shown for each odor presented (in columns), with neurons sorted by their time of peak firing relative to the onset of each odor (in rows). **c**, Accuracy of reconstructed time for each odor type was above chance levels. Plots show reconstructed time estimates obtained from each animal’s PSTH (correctly sorted for each odor; i.e., diagonal of panel b) and averaged across subjects. **d**, Accuracy of reconstructed time varied by the lag between the odor type used to train the model and the odor type in which time was reconstructed (e.g., when the model was trained on odor B trials, decoding time during odor B, C, D or E trials represented lags of 0, 1, 2, or 3, respectively). **e**, The sequential organization of firing fields extended across the full sequence of odors. Since the time between odor presentations varied across sequences and subjects (the task is self-paced), the PSTH (250ms gaussian) shows data from a single subject and the median onset of odors B, C, and D across sequences. **f**, Accuracy of reconstructed time across the sequence of odors (PSTH from panel e) was above chance levels. *, lag of 0 significantly different from other lags (Dunnett’s posthoc tests). L*, significant linear trend. Acc*, reconstructed time accuracy significantly above chance levels (determined by permutations).

### CA1 ensemble activity is organized in sequences of firing fields during sequence presentations

Consistent with previous reports^11–14^, hippocampal activity showed sequences of firing fields (“time cell” activity) during stimulus presentations (Fig 2b) as well as during intervals between stimulus presentations (Fig S1). Here we take advantage of our unique behavioral paradigm to extend these findings, by determining how this form of temporal coding interacts with stimulus identity and whether it extends across sequences of stimuli.

First, we found that these sequential patterns of activity were stimulus-specific and critical to sequence memory. For each odor type, strong temporal coding was observed when neurons were sorted by their peak firing latency relative to the onset of that odor (Fig 2c), whereas sorting the neurons according to the onset of another odor significantly reduced this temporal organization. More specifically, peri-event time histograms (PSTHs) involving the correct sorting were highly correlated across trials (PSTHs along the diagonal in Fig 2b; *r*_OdorA_ = 0.7797, *r*_OdorB_ = 0.7773, *r*_OdorC_ = 0.7632, and *r*_OdorD_ = 0.7631), and were significantly more correlated than when sorted by another odor (other PSTHs in same column; all one-way ANOVAs and Dunnett’s posthoc tests *P* values <0.05; Fig S2a). Similarly, our ability to accurately decode time based on the ensemble activity (accuracy of reconstructed time) was significantly higher when model training and decoding was performed on the same trial type (e.g., train and decode using odor A trials) than across trial types (e.g., train on odor A trials and decode B, C, or D trials; all one-way ANOVAs and Dunnett’s posthoc tests *P* values <0.05; Fig S2b). Importantly, reconstructed time accuracy was significantly higher before correct responses than incorrect responses (Kolmogorov-Smirnov D = 0.4692, *P* = 0.0011; tested on OutSeq trials for adequate sampling of correct and incorrect responses; −150ms to 100ms time window), suggesting this form of activity supports order memory decisions.

Second, we found that this form of coding systematically varied across (non-preferred) stimuli. In fact, consistent with the temporal context model^26, 27^, reconstructed time accuracy varied by lag (F_(6, 873)_ = 166.9, *P* <0.0001; Fig 2d). It was highest for lags of 0 (i.e., train and decode same odor type; all Dunnett’s posthoc tests *P* values <0.05) and showed a significant linear decline across lags of 1, 2, and 3 in the positive direction (e.g., train on B, decode C, D, or E; *F*_(1,327)_ = 12.93, *P* = 0.0004) and negative direction (e.g., train on D, decode C, B, or A; *F*_(1,327)_ = 12.20, *P* = 0.0005). It is also important to note that, despite the above differences, some of the temporal structure was also shared across trial types. In fact, reconstructed time accuracy was significantly above chance levels across all lags (Fig 2d; all one-sample *t*-tests *P* values <0.05; chance level determined by permutations), suggesting that this temporal organization is partially conserved across distinct experiences.

Finally, to determine if this temporal organization extended beyond individual stimuli or intervals, we examined the ensemble activity across the sequence of odors. Despite the variable time at which odors were presented (the task is self-paced so the time between odor presentations varied across sequences), the sequential organization of firing fields can be observed across the whole sequence (Fig 2e). Importantly, this activity provides significant information about time within sequences of events (Fig 2f), as reconstructed time accuracy was significantly higher than chance levels for each subject (*t*_(11)_ = 9.039, *t*_(6)_ = 6.859, *t*_(13)_ = 8.541, *t*_(11)_ = 6.092, *t*_(9)_ = 8.600; all *P* values <0.0005). These findings suggest that this form of sequential organization in CA1 ensemble activity can provide a task-critical temporal signal during stimulus presentations, during the interval separating stimulus presentations, as well as across sequences of stimuli.

### Differentiation of stimulus, temporal order, and trial outcome information varies within individual stimulus presentations

Next, we examined the temporal dynamics by which different types of task-critical information was represented within trials (wherein a trial corresponds to one odor presentation). Because nonspatial stimuli are less strongly represented in hippocampal neurons than spatial locations^28^, we used deep learning approaches to quantify the information represented in the ensemble activity. We began by identifying the underlying structure of ensemble activity at different moments within trials using a latent representation learning approach in which the model is not provided with any information about trial type. More specifically, we used an autoencoder, which can be viewed as a nonlinear counterpart to principal component analysis, to encode the spiking activity data into a two-dimensional latent space (by using only two neurons at the bottleneck of the neural network architecture; see methods). This approach allowed us to visualize a two-dimensional representation of the clustering of spike data at different time windows within trials, to which trial labels were subsequently added for posthoc analysis (see Fig 3a). The degree to which the resulting clusters mapped onto each type of trial information was quantified using a cross-validated *k*-nearest neighbor approach (*k* = 2) across animals, with high classification accuracy indicating high differentiation of a specific type of information (Fig 3b). For statistical comparisons, we determined the 95% confidence interval for chance-level classification accuracy for each type of information and time window using permutations. Accuracy values above the confidence intervals were considered statistically significant.

**Figure 3.**
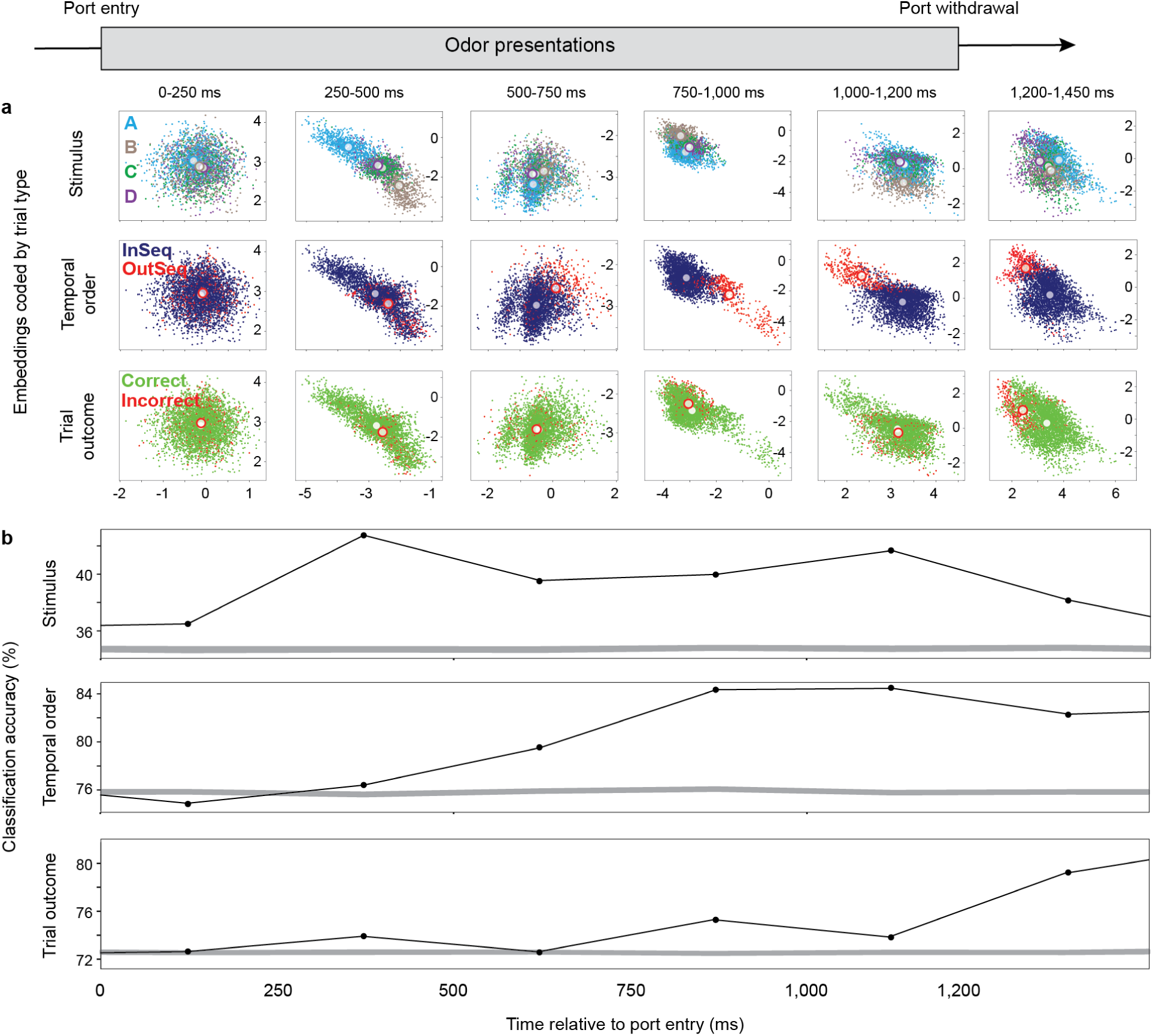
Stimulus, temporal order, and trial outcome information are simultaneously differentiated, but peak at different times, during stimulus presentation. A deep learning latent representation approach was used to identify the underlying structure of ensemble activity within 250ms bins, to which trial labels were subsequently applied. **a**, Differentiation of stimulus (odors), temporal order (InSeq or OutSeq), and trial outcome (correct or incorrect response) information in example subject. The model reduced the dimensionality of each animal’s ensemble activity to two dimensions, shown here for each 250ms window (each point represents a 100ms slice of spike train data, white circles represent cluster centroids). The axes refer to the two dimensions, the first and second latent variable, which correspond to the two nodes of the middle layer of the autoencoder (see Fig S4). The top row shows the model results color-coded by odor stimuli (correct InSeq trials only), the middle row by temporal order (correct InSeq and OutSeq trials), and the bottom row by trial outcome (all trials). **b**, Differentiation (classification accuracy) for each type of information relative to chance levels. Since two-dimensional embeddings were specific to each rat’s neuronal ensemble, classification accuracy was calculated using a cross-validated k-nearest neighbor approach (k = 2) and collapsed across subjects. Grey bands represent 95% confidence interval of chance-level classification accuracy at each time window (determined by permutations).

Despite the model not including trial-specific information, we identified patterns in the data that strongly differentiated information about stimulus (which odor was presented), temporal order (whether the odor was InSeq or OutSeq), and trial outcome (whether the animal correctly identified the trial or not). More importantly, this approach simultaneously captured the temporal dynamics of each type of information differentiation within trials. As expected, weak differentiation was observed immediately after port entry (0-250ms), that is before trial-specific information was available. Although above-chance odor differentiation was observed in that time period, which likely reflects prediction of the upcoming odor (see next section), the differentiation of temporal order or trial income information was at chance levels. The next time window (250-500ms) showed the strongest differentiation of odor information, but also significant differentiation of temporal order and trial outcome information. The differentiation of temporal order is consistent with previous single-cell analyses showing that a significant proportion of hippocampal neurons fire differentially on InSeq vs OutSeq trials^25^ (“sequence cells”), but the present analysis reveals that, at the ensemble level, this form of coding can be detected earlier in the trial. The trial outcome differentiation indicates the presence of a pattern in the ensemble activity that is predictive of whether the animal will respond correctly or not on that particular trial, which may reflect incorrect or disrupted representations of the predicted stimulus, currently presented stimulus, or InSeq/OutSeq status of the trial. Critically, these patterns of significant differentiation were observed while the animals’ behavior remained constant, as few port withdrawals occurred before 500ms^25^.

The key pattern in subsequent windows is that the odor differentiation was largely maintained (see next section), and that temporal order and trial outcome differentiation followed patterns consistent with the animals’ behavior. In fact, the third time window (500-750ms) is when most correct OutSeq responses were made (i.e., the animal was still in front of the port, but his nose was withdrawn). As expected, temporal order differentiation continued to rise in that time period (as most InSeq/OutSeq decisions were made), but little trial outcome differentiation was observed (as it was too early for the error signal to have been presented). Finally, the fourth (750-1,000ms) and sixth time (1,200-1,450ms) windows are when the majority of response feedback was provided for OutSeq and InSeq decisions, respectively, which resulted in an increase in trial outcome differentiation. More specifically, the fourth window (750-1,000ms) was when animals tended to receive feedback on whether their OutSeq decisions were correct (buzzer after error), and the sixth window (1,200-1,450ms) whether their InSeq decisions were correct (signal after correct, buzzer after incorrect). To summarize, our results show that ensemble activity simultaneously differentiated information about stimulus, temporal order, and trial outcome within individual stimulus presentations, and that the sequential organization of their respective peaks was consistent with the expected flow of task-critical information within trials.

### Sequential activation of upcoming stimuli within individual stimulus presentations

To follow up on the finding of sustained odor differentiation within trials, we then determined the content of the odor information represented. More specifically, we quantified the decoding of each odor representation at different moments within trials using a supervised model (a model that includes information about the stimulus presented on each trial). More specifically, we used a convolutional neural network (CNN) model with a tetrode-wise filtering architecture, which takes both spike data and LFP signals as input, and odor labels as its output. During training, the model was supplied trial-identified neural activity from the 150-400ms window and used multiple nonlinear hidden layers to create a map between input and output. After training, the hidden neural network layer provided a supervised latent representation of the data, which was used to decode odor representations across time windows. For statistical comparisons, we determined the 95% confidence intervals of the decoding probabilities in each time window (which follow a multinomial distribution). To be conservative, decoding probabilities with non-overlapping confidence intervals were considered significantly different.

Our results show that the representations of odors B, C, and D were sequentially activated within odor presentations. On odor B trials (Fig 4a,b), we found that the representation of B was significantly predicted before trial onset (−200ms to 50ms window). The representation of B was then maintained (significantly above that of other odors) until it peaked in the 200-450ms period. Of particular interest is the fact that, while still presented with odor B, the representation of upcoming items in the sequence (C and D) were activated in the subsequent time windows. In fact, the probability of B significantly declined as the probability of C and D increased (positive slope for “P_C_ – P_B_” difference across windows: *t*_6_ = 4.914, *P* = 0.00796; positive slope for “P_D_ – P_B_” difference: *t*_6_ = 5.069, *P* = 0.00357). Importantly, a similar pattern was observed on odor C trials (Fig 4c,d). In fact, not only was peak decoding of odor C occurring in the 200-450ms window and followed by the upcoming item in the sequence (odor D; positive slope for P_D_ – P_C_; *t*_6_ = 4.086, *P* = 0.0064), the representation of odor B was also activated before presentation of odor C (positive slope for P_C_ – P_B_; *t*_6_ = 9.078, *P* = 0.0008).

**Figure 4.**
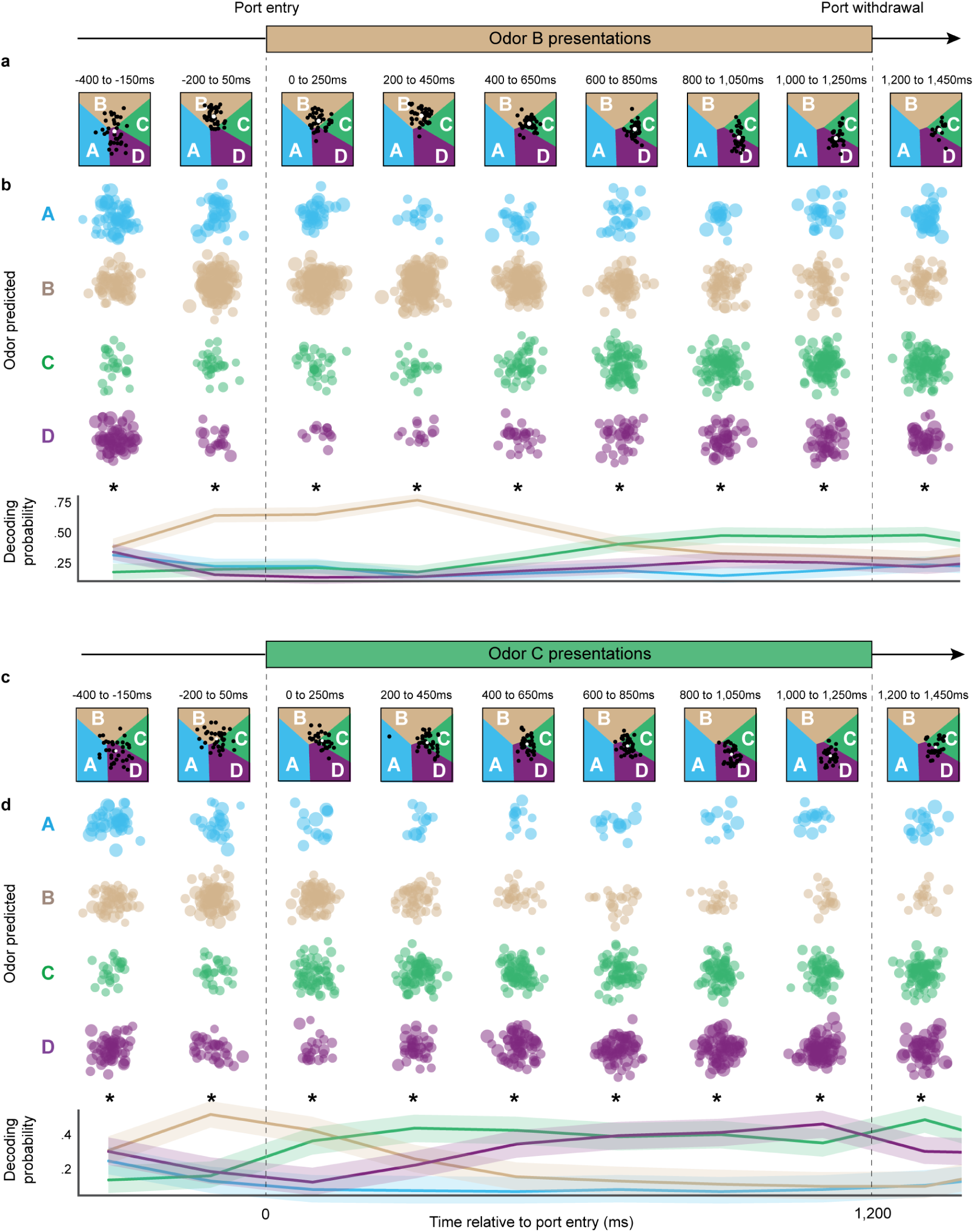
The neural representation of upcoming stimuli is sequentially activated during individual stimulus presentations. A deep learning neural decoding approach was used to quantify the decoding probability of each odor representation in 250ms bins. **a**, Decoding dynamics during odor B presentations in example subject. Regions of latent space are color coded according to the odor of highest probability, with boundaries indicating equal probability between two odors. Each point indicates the latent representation of a trial in that time window, with its position representing the decoded probabilities for each odor (cluster centroids shown as white circles). **b**, Group decoding dynamics during odor B presentations. *Top*, Since latent space coordinates were specific to each rat’s neuronal ensemble, data from each corresponding odor space and time window was collapsed across subjects. For each time window, data points (trials) were categorized and color-coded according to their decoded odor of highest probability, with the number of points in each cluster reflecting the decoded probability for each odor (equal number of points across time windows; size of points reflects decoded probability). The location of points within each cluster was randomly determined. *Bottom*, Summary plot of decoding probabilities over time (mean ±95% confidence intervals). **c**, Decoding dynamics during odor C presentations in same example subject. **d**, Group decoding dynamics during odor C presentations. *, significant chi-square test.

Similar sequential decoding peaks of B, C, and D were also observed on odor D trials (Fig S3), further supporting the consistency of the pattern. However, it is important to note that the representation of odor A was not strongly decoded throughout presentations of odors B, C, or D. This pattern is consistent with the fact that the first item of the sequence was always odor A (whereas any odor could be presented in subsequent sequence positions) and thus the discriminative part of the sequence started at the second sequence position. Finally, the pre-trial predictive coding above (decoding of B before odor B or C trials) was significantly stronger before correct than incorrect responses (Kolmogorov-Smirnov D = 0.3683, *P* = 0.0175; tested on OutSeq trials for adequate sampling of correct and incorrect responses), suggesting it is critically linked to order memory judgments. Collectively, these findings suggest that animals had developed a schema of the sequence, which was retrieved and played forward within each stimulus presentations to guide behavioral decisions.

### Representations of previous, current and future sequential stimuli are compressed within theta cycles

We then examined whether these sequential representations of nonspatial items could also be compressed within individual theta cycles, a phenomenon previously reported in place cell studies (theta sequences)^17, 18, 20^. During spatial navigation, theta sequences are thought to represent a segment of trajectory in space: within a single theta cycle, the animal’s preceding location tends to be represented in the descending phase, its current location at the trough, and its subsequent location in the ascending phase. Here, we tested the hypothesis that a similar temporal organization is observed for nonspatial sequences of stimuli that are normally separated by seconds (Fig 5a).

**Figure 5.**
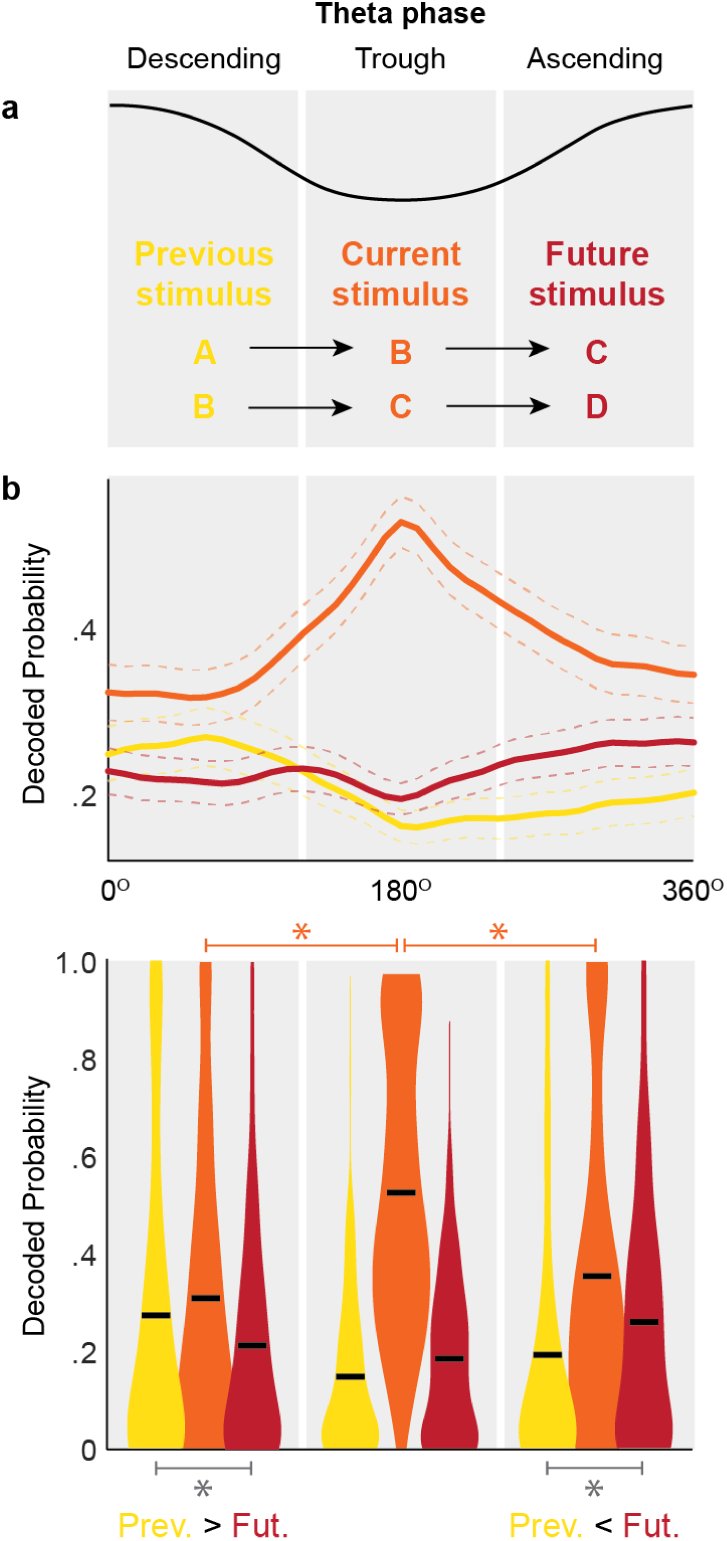
Representations of previous, current and future sequential stimuli are compressed within a theta cycle. A logistic regression model was used to quantify odor decoding within a theta cycle (the first cycle beginning 100ms after port entry). **a**, Hypothesized pattern if theta sequence compression previously reported in place cell studies extended to sequences of nonspatial stimuli: the currently presented stimulus (e.g., B) is predicted to be strongly represented during the trough of theta, the previous stimulus (e.g., A) during the ascending phase, and the future stimulus (e.g., C) during the descending phase. **b**, Decoding results from the three phases of theta support the hypothesized pattern. *Top*, Visualization of decoded probability for previous (yellow), current (orange), and future (red) stimuli during the theta cycle (collapsed across odor B and C trials, and across rats; mean ± 95% confidence interval). *Bottom*, Testing of hypothesized pattern: the descending phase showed significantly higher decoding of the previous stimulus than of the future stimulus, whereas the opposite pattern was found in the ascending phase, and decoding of the current stimulus was significantly stronger during the trough than either the ascending or the descending phase. *, significant paired t-test.

To do so, we used a simple multinomial logistic regression model to quantify odor decoding at faster timescales (while convolutional neural network models, like the one described in the previous section, provide more sensitive decoding, they require larger time windows that what is needed here). This analysis focused on the first theta cycle beginning 100ms after odor onset, to capture the period when hippocampal processing should be reflecting the currently presented odor. The model was trained using the spiking activity of the ensemble during the trough of that theta cycle (120-240 degrees; CA1 pyramidal layer theta) and was then used to decode odor information during the entire cycle (ascending phase, trough and descending phase; Fig 5a).

Consistent with evidence of theta sequences during spatial navigation, we found that information about previous, current, and future stimuli was differentially represented across the descending phase, trough, and ascending phase of theta (Fig 5b). More specifically, the currently presented stimulus was most strongly represented at the trough (*t*_DescendingVsTrough 294)_ = 10.2727, *P* <0.0001; *t*_TroughVsAscending (294)_ = 9.4514, *P* <0.0001), the descending phase showed significantly higher decoding of the previous stimulus than of the future stimulus (*t*_PreviousVsFuture (294)_ = 2.2098, *P* = 0.0189), but the opposite pattern was present in the ascending phase (future>previous; *t*_PreviousVsFuture (294)_ = 2.8302, *P* = 0.0025). These findings suggest that the representation of previous, current, and future nonspatial events, which are separated by several seconds in real time, can be compressed within theta cycles.

## DISCUSSION

In this study, we leveraged the use of novel behavioral and statistical approaches to identify new forms of sequential organization in hippocampal ensemble activity supporting the memory for sequences of events, a capacity known to critically depend on the hippocampus^**?**, 29, 30^. We report that sequential firing fields (time cell activity) were observed across a series of discrete nonspatial events distributed over several seconds, and that this temporal signal was linked with correct order memory judgments. We also found CA1 ensembles to be simultaneously and sequentially differentiating distinct types of trial-specific information within stimulus presentations, including the stimulus presented, its temporal order, and whether the animal correctly identified the trial type. In addition, we discovered that hippocampal ensembles replayed the corresponding sequential relationships among past, present, and future stimuli within individual stimulus presentations, and that these sequential representations could be compressed within a single theta cycle. Collectively, these results strongly suggest that encoding, preserving, and predicting event sequences is fundamental to hippocampal function. These findings are consistent with, and provide potential neuronal mechanisms for, the critical role of the hippocampus in temporally organizing our past experiences and future behavior.

Sequential firing fields (“time cells”) have previously been reported during individual stimulus presentations, inter-stimulus intervals, and across contiguous stimuli and responses^11–14^. Considerable evidence indicates this form of coding can provide a strong temporal signal within such task events^13, 14^. Here we offer, to our knowledge, the first direct evidence that hippocampal sequential firing fields provide significant temporal information across a sequence of discontinuous events unfolding over several seconds, and that this form of coding is crucial to remember the order of events. These findings lend support to theoretical models proposing that this form of coding supports our ability to organize our experiences in time^4, 26, 27, 31^, a characteristic feature of episodic memory^1–4^. In light of recent evidence that the lateral entorhinal cortex, a region with a strong anatomical and functional relationship with the hippocampus, provides a strong temporal signal across a longer timescale (several minutes)^32^, our results suggest that different aspects of the temporal context of episodic memories may be represented across medial temporal lobe structures.

Our findings support and extend recent evidence of theta-associated predictive and temporal coding properties in hippocampal ensembles. First, a previous study has reported theta-associated sequential representations that reflected the length of upcoming spatial trajectories^20^. These representations are thought to play a key role in planning and decision-making by maintaining an online representation of the planned trajectory toward the current goal. Importantly, the present study provides the first evidence that such predictive coding extends to nonspatial information, and that it is directly linked with correct sequence memory judgments. Notably, these findings are also compatible with the view that the hippocampus plays a key role in representing task-related schemas^33^, in this case the sequential relationships among those events, to guide behavioral decisions. Second, our findings are consistent with a recent study which used a conditional discrimination task with overlapping pairs of stimuli (a tone and an odor) to demonstrate theta phase precession for nonspatial information in hippocampal neurons^13^. In that study, the authors also used reconstructed time estimates to show that immediately preceding and upcoming time points (on the order of a few hundred ms) could be represented within a theta cycle. Our study extend these findings by suggesting a higher level of abstraction for nonspatial theta sequences — that the sequential relationships among preceding, current, and upcoming events (normally separated by seconds) can also be compressed within a theta cycle.

Our findings also raise a number of important questions to address in future studies. As we demonstrated, the sequence coding properties we reported were associated with correct order memory judgments. How can such representations in the hippocampus directly influence decisions and responses? It has been proposed that theta oscillations may play an essential role in allowing prospective hippocampal representations to facilitate decision-related processing in downstream structures^9, 20^. For instance, it is well established that neural activity in regions associated with planning behavior, including the prefrontal cortex and striatum, is significantly entrained to the hippocampal theta rhythm^34–36^. This proposal is consistent with recent work using this and other temporal memory tasks showing that, in addition to the hippocampus, the prefrontal cortex is also strongly engaged during task performance in both rodents and humans^23, 24, 37–40^. However, the specific prefrontal mechanisms remain to be demonstrated. Finally, it remains to be determined whether the novel forms of sequence coding reported here are present in other medial temporal lobe structures, and how these “online” theta-associated representations interact with “offline” sharp-wave-ripple-associated representations related to upcoming behavior or goals observed during reward consumption, quiescence, or sleep^15, 16^.

## METHODS

### Animals

Subjects were five male Long–Evans rats, weighing ∼350 g at the beginning of the experiment. They were individually housed and maintained on a 12 h light/dark cycle. Rats had ad libitum access to food, but access to water was limited to 2–10 min each day, depending on how much water they received as reward during behavioral training (3–6 ml). On weekends, rats received full access to water for ≥12 h to ensure adequate overall hydration. Hydration levels were monitored daily. All procedures were conducted in accordance with the Institutional Animal Care and Use Committee.

### Equipment and Stimuli

The apparatus consisted of a linear track (length, 150 cm; width, 9 cm), with walls angled outward (30°from vertical; height, 40 cm). An odor port, located on one end of the track, was equipped with photobeam sensors to precisely detect nose entries and was connected to an automated odor delivery system capable of repeated deliveries of multiple distinct odors. Two water ports were used for reward delivery: one located under the odor port, the other at the opposite end of the track. Timing boards (Plexon) and digital input/output boards (National Instruments) were used to measure response times and control the hardware. All aspects of the task were automated using custom Matlab scripts (MathWorks). A 96-channel Multichannel Acquisition Processor (MAP; Plexon) was used to interface with the hardware in real time and record the behavioral and electrophysiological data. Odor stimuli consisted of synthetic food extracts contained in glass jars (A, lemon; B, rum; C, anise; D, vanilla; E, banana) that were volatilized with desiccated, charcoal-filtered air (flow rate, 2 L/min). To prevent cross-contamination, separate Teflon tubing lines were used for each odor. These lines converged in a single channel at the bottom of the odor port. In addition, an air vacuum located at the top of the odor port provided constant negative pressure to quickly evacuate odor traces. Readings from a volatile organic compound detector confirmed that odors were cleared from the port 500–750 ms after odor delivery (inter-odor intervals were limited by software to ≥800 ms).

### Training

Naïve rats were initially trained to nosepoke and reliably hold their nose for 1.2 sec in the odor port for a water reward. Odor sequences of increasing length were then introduced in successive stages (A, AB, ABC, ABCD, and ABCDE) upon reaching behavioral criterion of 80% correct over three sessions per training stage. In each stage, rats were trained to correctly identify each presented item as either InSeq (by holding their nosepoke response for ∼ 1.2 sec to receive a water reward) or OutSeq (by withdrawing their nose before 1.2 sec to receive reward). There were two types of OutSeq items in the dataset, Repeats, in which an earlier item was presented a second time in the sequence (e.g., AB<u>A</u>DE), and Skips, in which an item was presented too early in the sequence (e.g., AB<u>D</u>DE, which skipped over item C). Although our previous work has revealed important differences in performance and neural activity on Repeats and Skips^22, 25^, this distinction was beyond the scope of the present analyses and not further discussed here. Note that OutSeq items could be presented in any sequence position except the first (i.e., sequences always began with odor A, though odor A could also be presented later in the sequence as a Repeat). After reaching criterion performance on the five-item sequence (> 80% correct on both InSeq and OutSeq items), rats underwent surgery for microdrive implantation.

### Surgery

Rats received a preoperative injection of the analgesic buprenorphine (0.02 mg/kg, s.c.) ∼10 min before induction of anesthesia. General anesthesia was induced using isoflurane (induction: 4%; maintenance: 1–2%) mixed with oxygen (800 ml/min). After being placed in the stereotaxic apparatus, rats were administered glycopyrrulate (0.5 mg/kg, s.c.) to help prevent respiratory difficulties. A protective ophthalmic ointment was then applied to their eyes and their scalp was locally anesthetized with marcaine (7.5 mg/ml, 0.5 ml, s.c.). Body temperature was monitored and maintained throughout surgery and a Ringer’s solution with 5% dextrose was periodically administered to maintain hydration (total volume of 5 ml, s.c.). The skull was exposed following a midline incision and adjustments were made to ensure the skull was level. Six support screws (four titanium, two stainless steel) and a ground screw (stainless steel; positioned over the cerebellum) were anchored to the skull. A piece of skull ∼3 mm in diameter (centered on coordinates: −4.0 mm AP, 3.5 mm ML) was removed over the left hippocampus. Quickly after the dura was carefully removed, the base of the microdrive was lowered onto the exposed cortex, the cavity was filled with Kwik-Sil (World Precision Instruments), the ground wire was connected, and the microdrive was secured to the support skull screws with dental cement. Each tetrode was then advanced ∼900*μm* into the brain. Finally, the incision was sutured and dressed with Neosporin and rats were returned to a clean cage, where they were monitored until they awoke from anesthesia. One day following surgery, rats were given an analgesic (flunixin, 2.5 mg/kg, s.c.) and Neosporin was reapplied to the incision site.

### Electrophysiological recordings

Each chronically implanted microdrive contained 20 independently drivable tetrodes. Following the surgical recovery period, tetrodes were slowly advanced over a period of ∼3 weeks while monitoring established electrophysiological signatures of the CA1 pyramidal cell layer (e.g., sharp waves, ripples, and theta amplitude). Voltage signals recorded from the tetrode tips were referenced to a ground screw positioned over the cerebellum, and filtered for single-unit activity (154 Hz to 8.8 kHz). The neural signals were then amplified (10,000–32,000x), digitized (40 kHz) and recorded to disk with the data acquisition system (MAP, Plexon). Action potentials from individual neurons were manually isolated off-line using a combination of standard waveform features across the four channels of each tetrode (Offline Sorter, Plexon). Proper isolation was verified using interspike interval distributions for each isolated unit (assuming a minimum refractory period of 1 ms) and cross-correlograms for each pair of simultaneously recorded units on the same tetrode. To confirm recording sites, current was passed through the electrodes before perfusion (0.9% PBS followed by 4% paraformaldehyde) to produce small marking lesions, which were subsequently localized on Nissl-stained tissue slices.

### Reconstructed time analyses

The objective of this approach was to quantify the degree of temporal information contained in sequential firing fields, by assessing the degree to which time can be accurately decoded from the ensemble activity. To do so, we adapted a memoryless Bayesian decoding model from a recent study examining temporal coding during sampling of olfactory and auditory stimuli^13^. Briefly, by assuming the ensemble spiking activity has a Poisson distribution and the spiking activity of each neuron is independent, we can use the spiking activity in a specific time window (actual time) to calculate the probability of time (reconstructed time) in Fig 2c as follows:

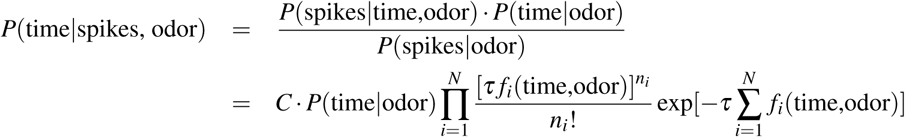

In Fig 2f, we use *P*(time, odor|spikes) instead. Here, *τ* is the length of the time window (50 ms with step size of 5ms in Fig 2c; 1500ms with step size of 150ms in Fig 2f), *f*_*i*_(time,odor) is the mean firing rate of the *i*-th unit in the time window, *n*_*i*_ is the number of spikes occurring in the time window, *odor* refers to the treatment of individual odors (odors ABCD in Fig 2c; odors ABCDE in Fig 2f), *C* is a normalization factor to make the probability distribution for each time window (actual time) sum up to 1, and assuming *time* is uniformly distributed given each odor, *P*(time|odor) is a constant since all InSeq trials have the same duration.

Unless otherwise noted, the model was trained on a balanced subset of sequence presentations in which all odors were InSeq, consecutively presented, and correctly identified. Plots of reconstructed time show, for each time window (actual time), the resulting probability distribution of reconstructed time averaged across subjects (red indicating max probability, blue indicating 0). Accuracy of reconstructed time was determined by quantifying the degree of relationship between actual time and reconstructed time estimates. To do so, for each trial, we calculated the correlation across rows and columns of the same index (one correlation value per row-column pair) and the mean of these correlation values represented the accuracy of reconstructed time on that specific trial. Chance levels were determined by calculating the mean reconstructed time accuracy across 1,000 random permutations of the time factor in the *f*_*i*_(time,odor) matrix. For the lag analysis, the model was tested on trial types not included in the training set. For lags of 1-3 (positive or negative) the model was trained using the proper sequence position of a given odor, but tested on the other sequence positions (e.g., training during B in ABCDE, but decoding during C, D, or E). For lags of 0, the model was tested on a subset of non-consecutive InSeq trials of comparable size. Comparisons between correct and incorrect trials focused on OutSeq trials because of the small number of errors observed on InSeq trials (i.e., the model was trained using InSeq trials, but tested using OutSeq trials). For each correct and incorrect OutSeq trial (collapsing across odor type), reconstructed time accuracy was calculated for a 250ms time window around port entry (−150ms to +100ms; a period preceding the identification of the currently presented odor during which the animals’ behavior was consistent across trials). Statistical comparisons between trial types were performed using ANOVAs and Dunnett’s posthoc tests with the exception of comparisons between correct and incorrect (OutSeq) trials, which used non-parametric Kolmogorov-Smirnov tests to account for potential non-normality in the distribution of values (though the same pattern of results was observed with parametric tests). Statistical comparisons with chance levels were performed using one-sample *t*-tests. All pairwise tests were two-tailed. Statistical significance was determined using *P*<0.05.

### Latent representation analyses

The objective of this approach was to visualize and quantify differences in the underlying structure of ensemble activity at different moments within individual stimulus presentations. To do so, we used an autoencoder, a nonlinear dimensionality reduction method based on neural networks^41, 42^, to identify a low-dimensional latent representation of the spike train data in 250ms time windows. Before applying the model, data from each stimulus presentation was divided into 250ms time windows (starting 2s before port entry, ending 2s after), and, for each time-window, a sliding window was used to extract 100ms sub-windows. The encoder portion of the model then projected the 100ms sub-windows of spike train data into a two-dimensional latent space by passing it through two layers with 500 nodes each and a “bottleneck” layer with 2 nodes, whereas the decoder portion of the model used the same layer architecture to project the latent space back into the original space to recover the input (see Fig S4). After training the model, we visualized the two-dimensional latent representation for each 250ms time window and color-coded each data point according to its trial type: the odor presented (stimulus), whether the odor was presented in or out of sequence (temporal order), and whether the rat performed the correct or incorrect response (trial outcome). Then, we used a *k*-nearest neighbors (*k*-NN) with *k* = 2 method to determine when the different types of trial-specific information were well separated in the latent space (70% of trials used for model training, 30% used for testing). More specifically, we predicted the stimulus, temporal order, and trial outcome label of each trial in the test set based on its top two closest neighbors in the training set. To focus the *k*-NN classification on the type of information of interest, the *k*-NN classification of stimulus and temporal order information only included correct trials (to focus on trial-relevant representations) and odors B, C, and D (to focus on odors with comparable discriminability; including odor A, which was strongly differentiated by the model, further enhanced classification accuracy). Similarly, stimulus classification only included for InSeq trials. For statistical comparisons, we determined the 95% confidence interval for chance-level classification accuracy for each type of information and time window using 100 permutations. For each permutation, we randomly shuffled the labels corresponding to stimulus, temporal order, and trial outcome and repeated *k*-nearest neighbors classification. Accuracy values above the confidence intervals were considered statistically significant.

### Neural decoding analyses

The objective of this approach was to build a powerful predictive model for decoding the stimulus information represented in the neural activity data using a supervised learning method. More specifically, we used a convolutional neural network (CNN), an approach most commonly used for image classification^43^ but that can also be used for sequential data^44^. For our analysis, the CNN took both spike data and LFP signals from each tetrode as input, and odor labels as its output. The convolution was performed on each tetrode separately (Fig S5a). More specifically, for each implanted tetrode, the continuous LFP signal and spike trains from each unit were combined into one multivariate time series. Filters (small matrices) were then convolved on these time series to produce convolution features. Time averaged features from different tetrodes were concatenated together and fed into subsequent hidden layers with drop-out^45^ and, finally to the output layer prediction. The model was trained using the 150-400ms time window and, as shown in Fig S5b, produced higher decoding accuracy compared to the benchmark multinomial logistic regression model (10-fold cross-validation). Moreover, by using the last hidden layer as a supervised latent space, our model allows for nonlinear projection of the original data onto a low-dimensional space for visualizing and quantifying the decoding probability of each odor over time. Note that in contrast to the autoencoder model used for the latent representation learning analyses, this latent space was obtained in a supervised manner by taking the labels (as well as the neural signals) into account. For each trial, the decoding model was then applied to each 250ms time window (ranging from −400ms to 1,450ms, relative to port entry) to obtain the corresponding hidden layer vectors. After the latent space was divided into subregions associated with each odor (according to the odor of highest probability), the latent space vectors for each trial could be visualized as a point that moves around in the latent space across the different time windows. For easier visualization (see Fig 4a,c), we have further reduced the dimension of the latent space to two using the top two principal components and linearized the boundaries between odors using a simple multinomial logistic regression model. To aggregate decoding results across subjects, for each time window, we compiled the number of points (trials) in each odor subregion of each subject’s latent space. As a result, for each time window, we obtained four clusters of points representing the aggregate decoding probability of odors A, B, C and D (the total number of points remains constant across windows). For statistical comparisons, we determined the 95% confidence intervals of the decoding probabilities in each time window, which follow a multinomial distribution. To be conservative, decoding probabilities with non-overlapping confidence intervals were considered significantly different.

### Theta sequence decoding analyses

The objective of this approach was to determine whether the sequential activation of odor representations can be detected in a compressed form within a single theta cycle. To do so, we used a LASSO logistic regression model^46^ that imposes *L*_1_ penalty on the regression parameters. Although this model yields weaker odor decoding than the CNN model (Fig S5b), it can be used at the faster timescale necessary to decode within a theta cycle (the CNN model requires larger time bins). Notably, the logistic regression model shows a similar pattern of sequential activation of upcoming stimuli within individual presentations as the CNN model, though, as expected, with more variability (Fig S6). For each trial (−2s to 2s, relative to port entry), the LFP signal from each tetrode was smoothed with a Butterworth filter between 4Hz and 7Hz (the main frequency range of theta we observed during odor sampling^25^) and the Hilbert transformation was then used to calculate the theta phase at each time point. The model was trained using the firing rate of each neuron during the trough (120-240 degrees; CA1 pyramidal layer theta) of the first theta cycle beginning 100ms after port entry. Trials in which the amplitude of that theta cycle was very weak (lowest 20th percentile) were not included in the analysis. Since not every neuron’s activity was associated with odor presentations, the LASSO model automatically eliminated neurons not significantly contribution to the odor decoding. The amount of penalty was set using a 10-fold cross-validation with stratification because the numbers of trials for different odors were unequal^47^. To apply the decoding model to the entire theta cycle, we used a sliding window (120 degrees in width) to calculate the neuron firing rates in 10 degree increments and determine the decoding probability of each odor. Paired *t*-tests were used to test the hypothesized pattern (Fig 5a) of decoding for previous, current, and future stimuli across the three phases of that theta cycle (descending phase, trough, and ascending phase). To maximize statistical power, decoding results were combined across odor B and C trials. Statistical significance was determined using *P*<0.05.

## ACKNOWLEDGEMENTS

This work was supported by NIH grants R01-MH115697 and R01-DC017687, NSF awards IOS-1150292 and BCS-1439267, and Whitehall Foundation award 2010-05-84.

## SUPPLEMENTAL INFORMATION

**Figure S1.**
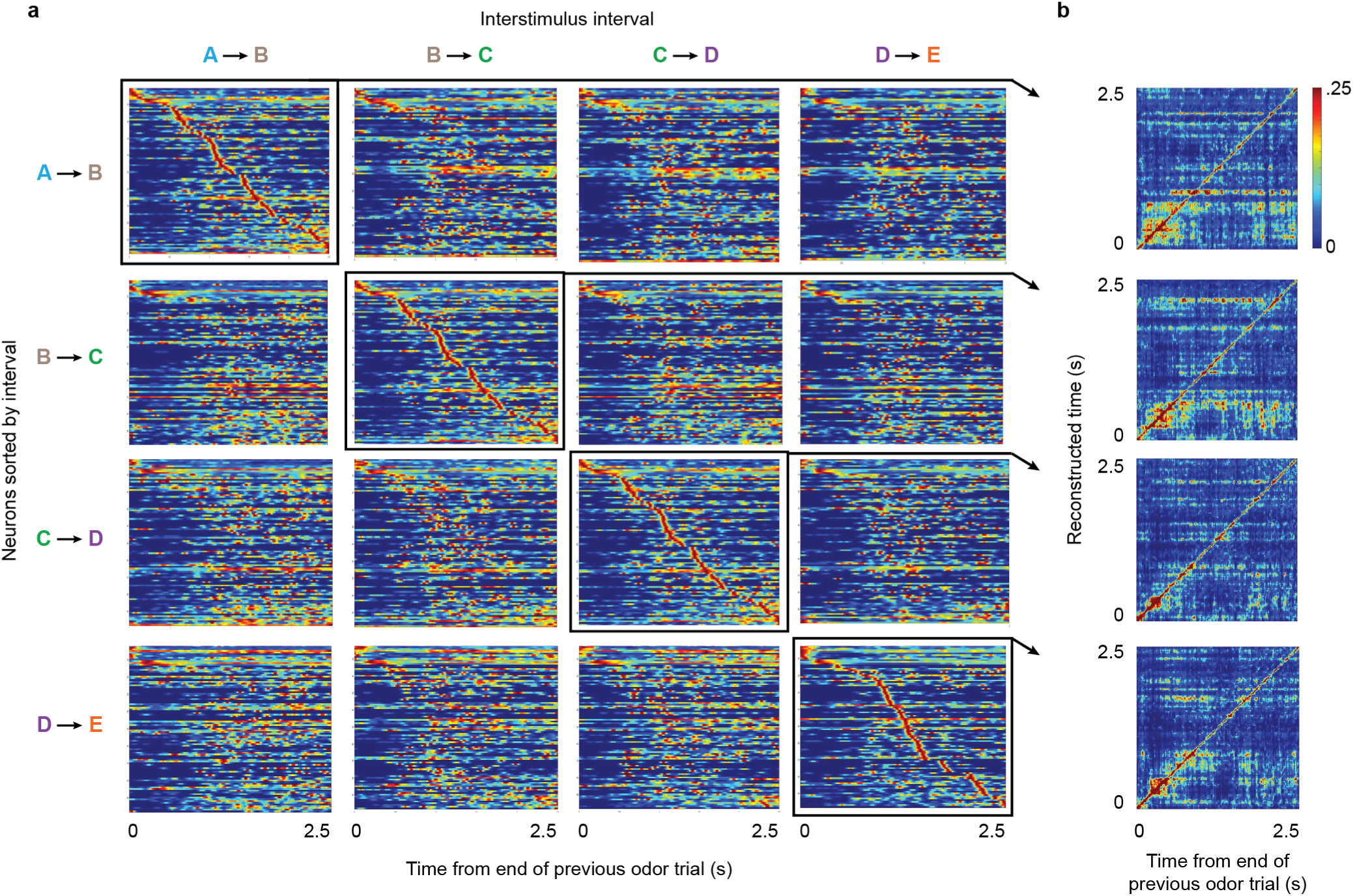
Example ensemble activity during inter-stimulus intervals. **a**, Since the duration of inter-stimulus intervals varied across animals, ensemble activity is shown from a representative subject (0s corresponds to reward delivery at end of previous odor presentation; plots are terminated at 2.5s but some intervals were of longer duration). Each peri-stimulus time histogram (PSTH) shows the normalized firing rate of all active neurons during a specific inter-stimulus interval (red indicates the maximum firing rate of each neuron, and blue indicates 0 Hz; 98 active neurons for each interval, except for B-C interval which had 99 active neurons). To visualize how sequential firing fields varied across intervals, PSTHs are shown for each interval (in columns), with neurons sorted by their time of peak firing relative to other intervals (in rows). **b**, Accuracy of decoded time during correctly sorted inter-stimulus intervals (i.e., diagonal of panel a).

**Figure S2.**
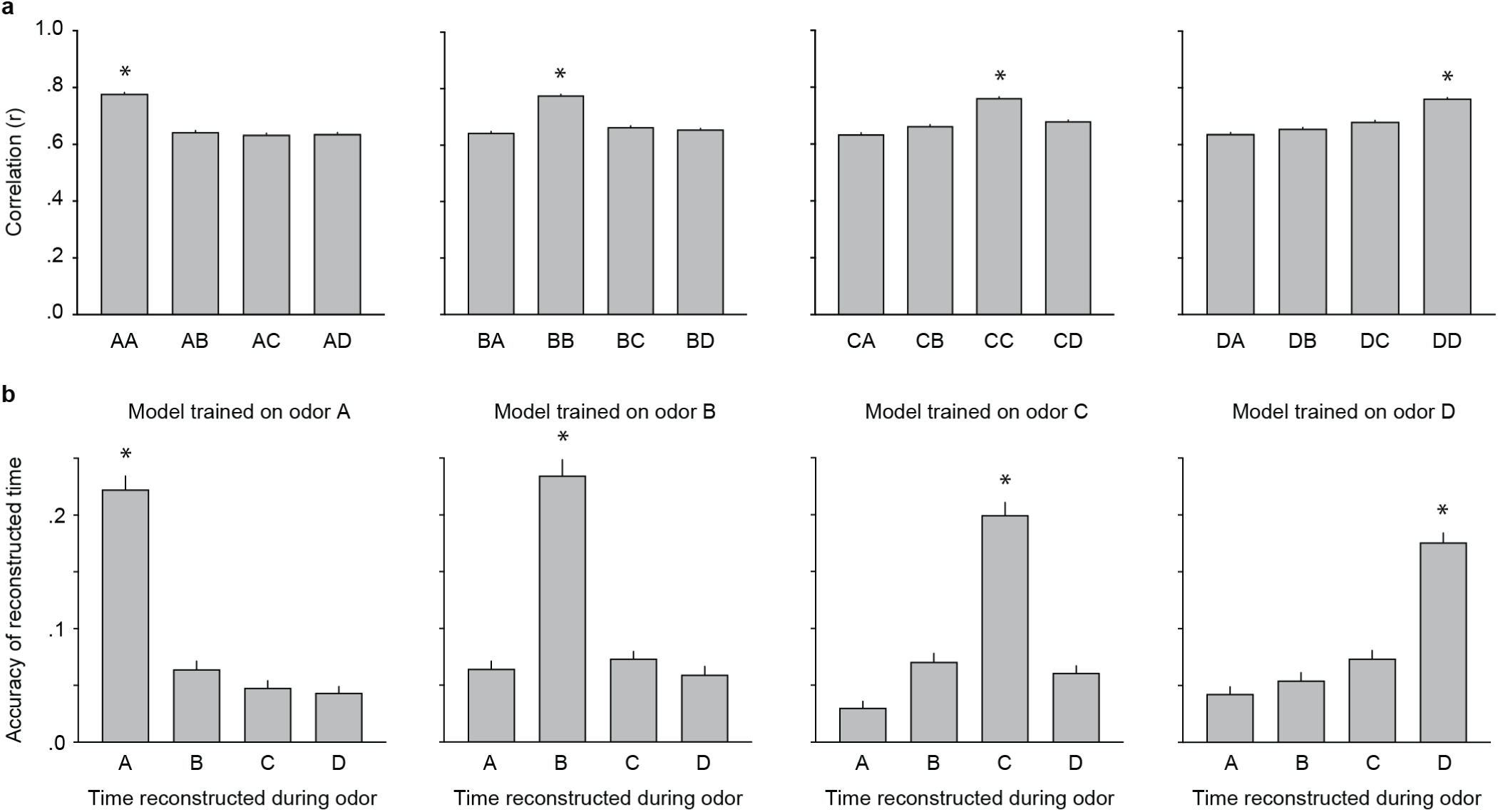
Stimulus-specificity of sequential firing fields. **a**, Mean correlation of PSTHs across trials of each odor type. An ANOVA showed that the pairwise correlations between each odor A trial and each other A, B, C, or D trial were significantly different across trial types (F_(3, 52)_ = 336.9, *P* <0.0001), with correlations across trials of the same type being significantly higher than the others (AvsA was significantly higher than AvsB, AvsC, or AvsD; Dunnett’s posthoc tests *P* values <0.05). The same pattern of results was obtained when the corresponding analyses were performed for odor B trials (B vs A,B,C, or D: *F*_(3, 52)_ = 334.4, P <0.0001; BvsB significantly higher than others), odor C trials (C vs A,B,C, or D: *F*_(3, 52)_ = 258.2, P <0.0001; CvsC significantly higher than others), and odor D trials (D vs A, B, C, or D: *F*_(3, 52)_ = 237.9, *P* <0.0001; DvsD significantly higher than others). **b**, Analysis of reconstructed time across trial types. An ANOVA showed that, when the model was trained using odor A trials, its reconstructed time accuracy significantly varied when applied to odor A, B, C, or D trials (*F*_(3, 216)_ = 113.1, *P* <0.0001), with the highest accuracy observed when decoding the trial type as in model training (Dunnett’s posthoc tests *P* values <0.0001). Again, a similar pattern of results was obtained when the model was trained using odor B trials (*F*_(3, 216)_ = 85.33, *P* <0.0001; Dunnett’s posthoc tests *P* values <0.0001), odor C trials (*F*_(3, 216)_ = 88.77, *P* <0.0001; Dunnett’s posthoc tests *P* values <0.0001), and odor D trials (*F*_(3, 216)_ = 71.27, *P* <0.0001; Dunnett’s posthoc tests *P* values <0.0001). *, decoding of same odor type as model training significantly higher than other odor types (Dunnett’s posthoc tests).

**Figure S3.**
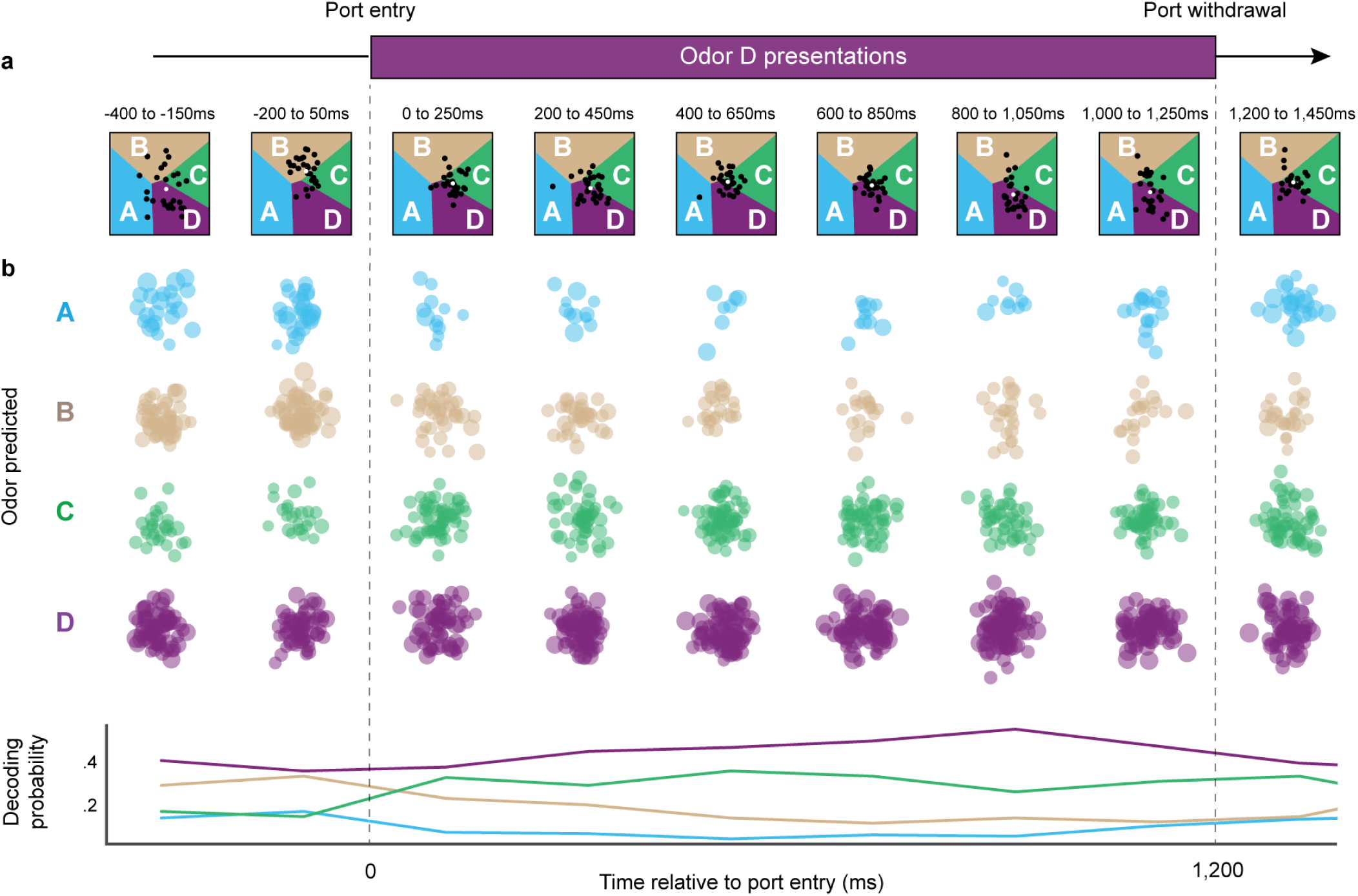
Sequential activation of upcoming stimuli during odor D presentations. **a**, Decoding dynamics during odor D presentations in example subject. **b**, Group decoding dynamics during odor D presentations.

**Figure S4.**
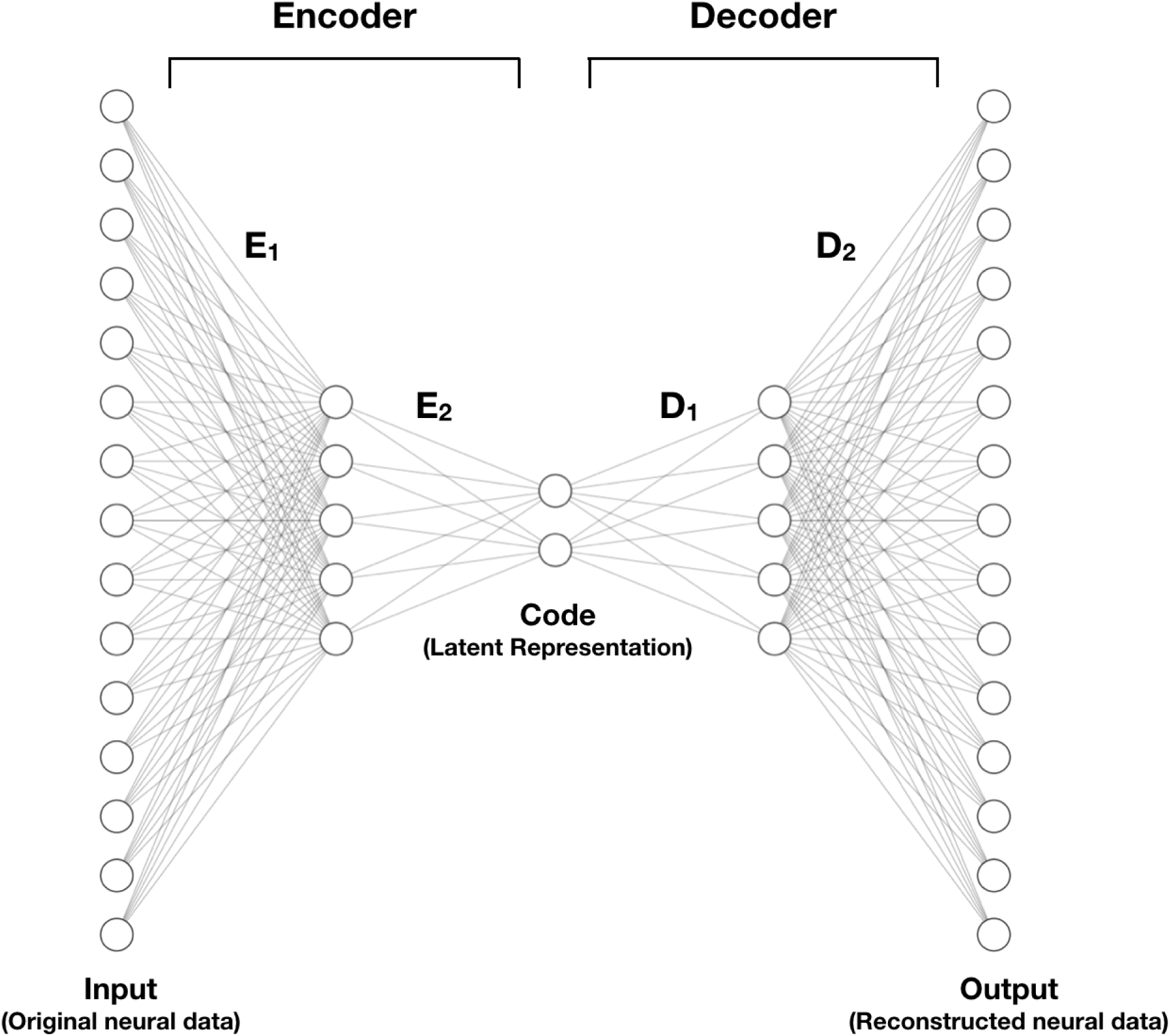
Illustration of the autoencoder used for latent representation analyses. The encoder and decoder components consist of “layers”, where each layer includes a number of nodes. Node *i* of layer *l* is defined as 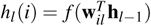, where *f* is some pointwise function, **w**_*il*_ is a vector of learnable parameters, and **h**_*l*−1_ represents the output of the previous layer. The input layer (observed data) is denoted as **h**_0_. The model was trained to minimize the difference between the original input data and the reconstructed data. To accomplish this, we estimated the parameters **w**_*il*_, for all *i* and *l*, using stochastic gradient descent^48^ with momentum^49^ to minimize the mean squared error between the input and output. The encoder portion projected the spike train data into a two-dimensional latent space by passing it through two layers with 500 nodes each and a “bottleneck” layer with 2 nodes. The decoder portion projected the latent space back into the original space by passing the output from the bottleneck layer through two layers with 500 nodes each and finally to an output layer whose dimensionality was the same as the input. We used a rectified non-linear unit for the pointwise function^50, 51^, *f*, for all layers except the bottleneck layer, where we used a linear function.

**Figure S5.**
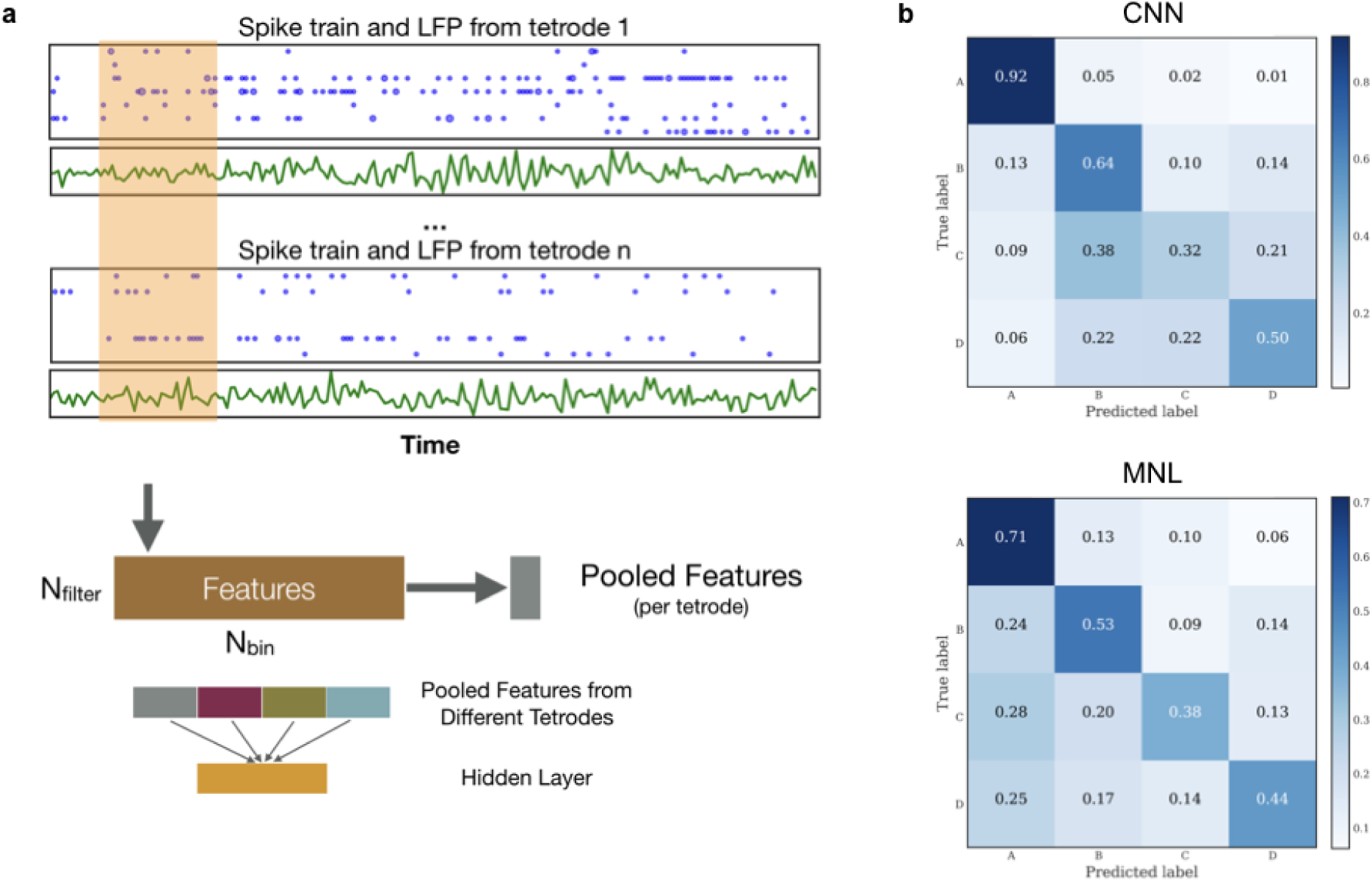
Illustration of the convolutional neural network (CNN) used for the neural decoding analyses, and comparison to a multinomial logistic (MNL) regression model. **a**, Neural activity from each trial (−2s to 2s relative to port entry) was segmented into 250ms time windows. The model mapped the LFP and spike train data within the time window to a hidden layer vector and then made a prediction based on that vector. More specifically, the neural network performed convolution on each tetrode separately. The LFP signal and spike trains obtained from each tetrode were combined into one multivariate time series passing through filters to produce convolution features. Next, time-averaged features from all the tetrodes were concatenated together and passed into a hidden layer before output prediction. For each trial, the 250ms time window starting from 0.15s after port entry was used to train the decoding model. For training the neural network, we used a variant of stochastic gradient descent algorithm^52^ and followed the early stopping rule. **b**, Confusion matrices for the CNN (*top*) and MNL with LASSO penalty (*bottom*). Each confusion matrix shows the proportions of correct and incorrect model classifications for each odor. The CNN and MNL models were both trained on the same 150-400ms window for all the rats with 10-fold cross-validation. The results shown were calculated using model predictions on the validation folds. The overall classification accuracy across all four odors and five rats of the CNN model was 64.5% while the LASSO logistic regression model had 54.6%. The CNN had higher classification accuracy than the MNL model for all the odors except odor C. More importantly, the CNN model was able to better preserve the sequential pattern of odors whereas the MNL model mis-classified many trials to odor A (as seen in the first column on the right).

**Figure S6.**
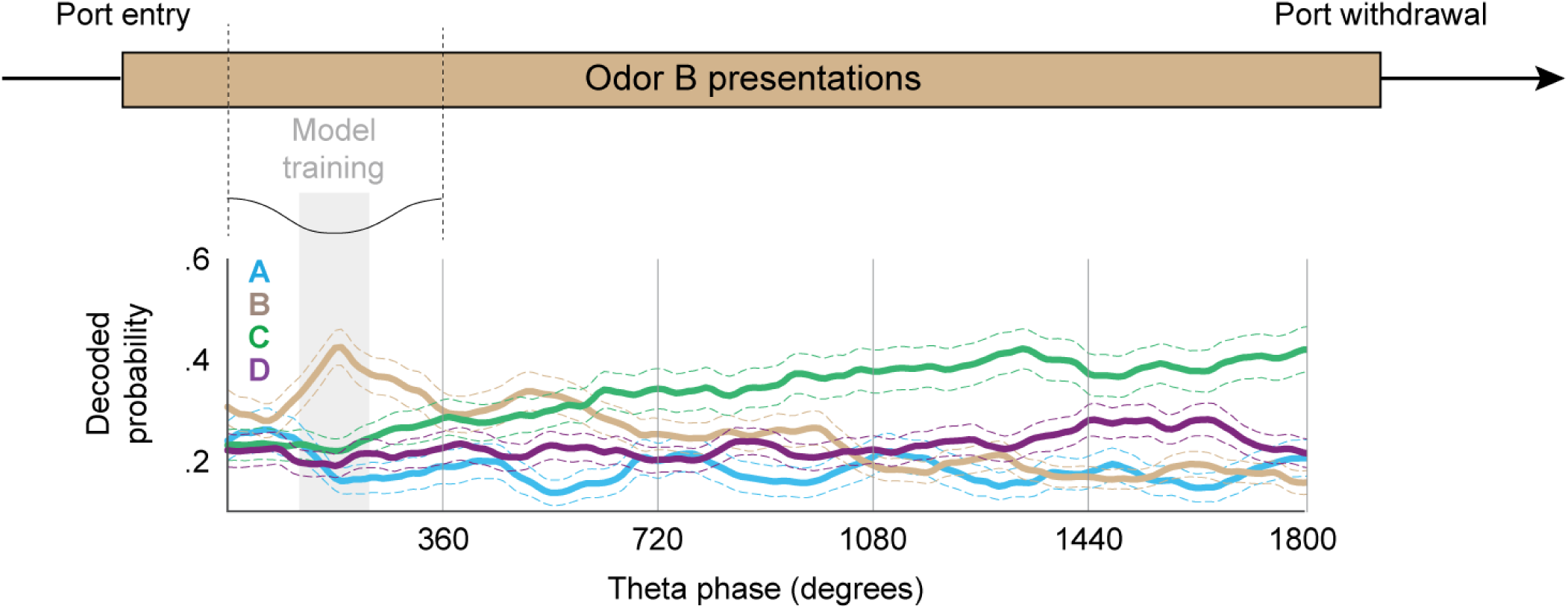
Decoding of sequential activation of upcoming stimuli using the multinomial logistic regression model. Despite overall weaker odor decoding than the CNN model, the logistic regression model shows a similar pattern of sequential activation of upcoming stimuli (compare with Fig 4b).

## References

1. Clewett, D., DuBrow, S. & Davachi, L. Transcending time in the brain: How event memories are constructed from experience. Hippocampus 29, 162–183 (2019).

2. Ranganath, C. & Hsieh, L. T. The hippocampus: A special place for time. AnnalsNYAcadSci 1369, 93–110 (2016).

3. Allen, T. A. & Fortin, N. J. The evolution of episodic memory. PNAS 110, 10379–10386 (2013).

4. Eichenbaum, H. Time cells in the hippocampus: A new dimension for mapping memories. Nature Reviews Neuroscience 15, 732–744 (2014).

5. Schacter, D. L. et al. The Future of Memory: Remembering, Imagining, and the Brain. Neuron 76, 677–694 (2012).

6. Redish, A. D. Vicarious trial and error. Nature Reviews Neuroscience 17, 147–159 (2016).

7. Lisman, J. E. Relating Hippocampal Circuitry Viewpoint to Function: Recall of Memory Sequences by Reciprocal Dentate–CA3 Interactions. Neuron 22, 233–242 (1999).

8. Levy, W. B. A sequence predicting CA3 is a flexible associator that learns and uses context to solve hippocampal-like tasks. Hippocampus 6, 579–590 (1996).

9. Buzsáki, G. & Tingley, D. Space and Time: The Hippocampus as a Sequence Generator. Trends in Cognitive Sciences 22, 853–869 (2018).

10. Lisman, J. & Redish, A. D. Prediction, sequences and the hippocampus. Phil. Trans. R. Soc. B 364, 1193–1201 (2009).

11. Pastalkova, E., Itskov, V., Amarasingham, A. & Buzsaki, G. Internally Generated Cell Assembly Sequences in the Rat Hippocampus. Science 321, 1322–1327 (2008).

12. MacDonald, C. J., Lepage, K. Q., Eden, U. T. & Eichenbaum, H. Hippocampal “time cells” bridge the gap in memory for discontiguous events. Neuron 71, 737–749 (2011).

13. Terada, S., Sakurai, Y., Nakahara, H. & Fujisawa, S. Temporal and Rate Coding for Discrete Event Sequences in the Hippocampus. Neuron 94, 1248–1262 (2017).

14. Mau, W. et al. The Same Hippocampal CA1 Population Simultaneously Codes Temporal Information over Multiple Timescales. Current Biology 28, 1499–1508 (2018).

15. Foster, D. J. Replay Comes of Age. Annual Review of Neuroscience 40, 581–602 (2017).

16. Joo, H. R. & Frank, L. M. The hippocampal sharp wave–ripple in memory retrieval for immediate use and consolidation. Nature Reviews Neuroscience 19, 744–757 (2018).

17. Skaggs, W. E., McNaughton, B. L., Wilson, M. A. & Barnes, C. A. Theta Phase Precession in Hippocampal Neuronal Populations and the Compression of Temporal Sequences. Hippocampus 6, 149–172 (1996).

18. Foster, D. J. & Wilson, M. A. Hippocampal Theta Sequences. Hippocampus 17, 1093–1099 (2007).

19. Dragoi, G. & Buzsáki, G. Temporal Encoding of Place Sequences by Hippocampal Cell Assemblies. Neuron 50, 145–157 (2006).

20. Wikenheiser, A. M. & Redish, A. D. Hippocampal theta sequences reflect current goals. Nature Neuroscience 18, 289–294 (2015).

21. Johnson, A. & Redish, A. D. Neural Ensembles in CA3 Transiently Encode Paths Forward of the Animal at a Decision Point. Journal of Neuroscience 27, 12176–12189 (2007).

22. Allen, T. A., Morris, A. M., Mattfeld, A. T., Stark, C. E. & Fortin, N. J. A sequence of events model of episodic memory shows parallels in rats and humans. Hippocampus 24, 1178–1188 (2014).

23. Boucquey, V., Allen, T., Huffman, D., Fortin, N. & Stark, C. Memory for sequences of events shows bilateral hippocampal and medial prefrontal cortical activity in humans (under review). In Society for Neuroscience Abstracts (Washington, DC, 2014).

24. Fortin, N. et al. Distinct contributions of hippocampal, prefrontal, perirhinal and nucleus reuniens regions to the memory for sequences of events. In Society for Neuroscience Abstracts (San Diego, CA, 2016).

25. Allen, T. A., Salz, D. M., McKenzie, S. & Fortin, N. J. Nonspatial sequence coding in CA1 neurons. Journal of Neuroscience 36, 1547–1563 (2016).

26. Howard, M. W. & Kahana, M. J. A distributed representation of temporal context. Journal of Mathematical Psychology 46, 269–299 (2002).

27. Shankar, K. H. & Howard, M. W. Timing using temporal context. Brain Research 1365, 3–17 (2010).

28. Manns, J. R., Howard, M. W. & Eichenbaum, H. Gradual Changes in Hippocampal Activity Support Remembering the Order of Events. Neuron 56, 530–540 (2007).

29. Fortin, N. J., Agster, K. L. & Eichenbaum, H. B. Critical role of the hippocampus in memory for sequences of events. Nature Neuroscience 5, 458–462 (2002).

30. Kesner, R. P., Gilbert, P. E. & Barua, L. A. The role of the hippocampus in memory for the temporal order of a sequence of odors. Behavioral Neuroscience 116, 286–290 (2002).

31. Shankar, K. H. & Howard, M. W. A Scale-Invariant Internal Representation of Time. Neural Computation 24, 134–193 (2012).

32. Tsao, A. et al. Integrating time from experience in the lateral entorhinal cortex. Nature (2018).

33. McKenzie, S. et al. Representation of memories in the cortical–hippocampal system: Results from the application of population similarity analyses. Neurobiology of Learning and Memory 134, 178–191 (2016).

34. Van Der Meer, M. A. A. & Redish, A. D. Theta phase precession in rat ventral striatum links place and reward information. Journal of Neuroscience 31, 2843–2854 (2011).

35. Jones, M. W. & Wilson, M. A. Phase precession of medial prefrontal cortical activity relative to the hippocampal theta rhythm. Hippocampus 15, 867–873 (2005).

36. Hyman, J. M., Zilli, E. A., Paley, A. M. & Hasselmo, M. E. Medial prefrontal cortex cells show dynamic modulation with the hippocampal theta rhythm dependent on behavior. Hippocampus 15, 739–749 (2005).

37. Allen, M., Leila Lesyshyn, R. A. J. O. S., Allen, T. A. & Fortin, N. J. The hippocampus, prefrontal cortex, and perirhinal cortex are critical to incidental order memory. Behavioural Brain Research https://doi.org/10.1016/j.bbr.2019.112215 (2019).

38. Chiba, A. A., Kesner, R. P. & Reynolds, A. M. Memory for spatial location as a function of temporal lag in rats: role of hippocampus and medial prefrontal cortex. Behavioral and Neural Biology 61, 123–131 (1994).

39. Jenkins, L. J. & Ranganath, C. Prefrontal and Medial Temporal Lobe Activity at Encoding Predicts Temporal Context Memory. Journal of Neuroscience 30, 15558–15565 (2010).

40. Schuck, N. W. & Niv, Y. Sequential replay of nonspatial task states in the human hippocampus. Science (New York, N.Y.) 364, eaaw5181 (2019).

41. DeMers, D. & Cottrell, G. W. Non-linear dimensionality reduction. In Advances in neural information processing systems, 580–587 (1993).

42. Hinton, G. E. & Salakhutdinov, R. R. Reducing the dimensionality of data with neural networks. science 313, 504–507 (2006).

43. LeCun, Y., Bottou, L., Bengio, Y., Haffner, P. et al. Gradient-based learning applied to document recognition. Proceedings of the IEEE 86, 2278–2324 (1998).

44. Goodfellow, I., Bengio, Y. & Courville, A. Deep learning (MIT press, 2016).

45. Baldi, P. & Sadowski, P. J. Understanding dropout. In Advances in neural information processing systems, 2814–2822 (2013).

46. Tibshirani, R. Regression shrinkage and selection via the lasso. Journal of the Royal Statistical Society: Series B (Methodological) 58, 267–288 (1996).

47. Friedman, J., Hastie, T. & Tibshirani, R. The elements of statistical learning, vol. 1 (Springer series in statistics New York, 2001).

48. Rumelhart, D. E., Hinton, G. E., Williams, R. J. et al. Learning representations by back-propagating errors. Cognitive modeling 5, 1 (1988).

49. Sutskever, I., Martens, J., Dahl, G. & Hinton, G. On the importance of initialization and momentum in deep learning. In International conference on machine learning, 1139–1147 (2013).

50. Nair, V. & Hinton, G. E. Rectified linear units improve restricted boltzmann machines. In Proceedings of the 27th international conference on machine learning (ICML-10), 807–814 (2010).

51. Jarrett, K., Kavukcuoglu, K., Ranzato, M. & LeCun, Y. What is the best multi-stage architecture for object recognition? In 2009 IEEE 12th international conference on computer vision, 2146–2153 (IEEE, 2009).

52. Kingma, D. P. & Ba, J. Adam: A method for stochastic optimization. arXiv preprint 1412.6980 (2014).

